# Out-of-sight or out-of-site? Forays and detection probabilities in single-season occupancy models

**DOI:** 10.1101/2021.05.18.444729

**Authors:** André Desrochers, Pierre Blanchette, Marc J. Mazerolle

## Abstract

Occupancy models have become popular in wildlife survey analyses because they account for the frequent failure to detect individuals of targeted species. Those individuals sometimes move outside sampling sites, i.e. exhibit temporary emigration. In such cases, occupancy models may become difficult to interpret or even misleading either at the species or the individual level, because they confound presence at the site, availability for detection given presence, and actual detection by the observer. We quantified the probabilities of these three components with spruce grouse (*Falcipennis canadensis*) in southern Quebec, Canada. We conducted call-response surveys of 24 grouse monitored by radio-telemetry. We defined sites empirically as circular areas of 83 m radius centered on the observer, corresponding to the maximum detection distance obtained. Based on telemetry locations, grouse were present at the site during 42 % of the surveys. Six stationary grouse were present during surveys, but were never detected. Thus, only individuals that moved in the presence of the observer (89 %) were considered available for detection. Individuals available for detection were detected in 51 % of the cases. We simulated detection histories and built single-season occupancy models, based on the empirical relationship between detection probability and the distance measured between observers and grouse. When temporary emigration was ignored, site occupancy was *ψ* = 0.89, and the associated probability of detection was *p* = 0.23. When instances of temporary emigration were dropped, estimates were *ψ* = 0.88 and *p* = 0.41. Using only grouse available for detection, estimates were *ψ* = 0.87 and *p* = 0.42. Disentangling the components of detection probabilities had little impact on occupancy estimates, but showed a major effect of temporary emigration on estimated detection probabilities.

## Introduction

In most bird species, the detection probability of individuals is rarely perfect during a survey season. It may be affected by factors such as observer experience, weather, vegetation structure, time of the season, time of day, and behaviour. The detection process is the product of at least three components, each with probability < 1: presence at a sampling site during the survey (psi, *ψ*), availability for detection (e.g., bird in view or calling) given presence (*p_a_*), actual detection given presence and availability (*p_d_*; (Kéry and Schmidt, 2008).

Occupancy, i.e., the probability that an individual or a species is present at a given location during a given period of time (MacKenzie et al., 2006), is a state variable of interest in assessing habitat relationships and understanding population responses to management practices or environmental change. Failure to detect individuals not only leads to underestimation of true occupancy, but in cases where detection is dependent on habitat or sampling conditions, it may lead to erroneous inference when examined with conventional methods, e.g. generalized linear models (MacKenzie et al., 2006; Mazerolle et al., 2005). Consequently, statisticians have developed analytical methods to estimate species occupancy while accounting for the probability of detection given presence, to improve inferences from large-scale occurrence patterns and dynamics (MacKenzie et al., 2002, 2006).

Based on repeated surveys at a fixed array of sites, single-season occupancy models enable the estimation of both the probability that a site is occupied by the focal individual or species (occupancy, *ψ*) and the probability of detecting it at time *t* (*p_t_*), given its presence at the site. Single-season occupancy models typically assume closure – a site remains occupied or unoccupied between the first and last visits conducted at the site during the season. However, closure, as well as occupancy and detection probabilities, depend on establishing temporal and spatial scales relevant to the species of interest. In cases where the entire sites of interest are sampled (e.g., amphibians in a set of small ponds), closure during the breeding period is a reasonable assumption, and occupancy estimates will be straightforward to interpret. However, closure may be unrealistic if sampling sites represent a small or unknown proportion of either the area occupied by the targeted species, or the home range of the targeted individual. In the latter situation, the interpretation of occupancy model results is difficult, and a looser concept, “use”, is recommended, rather than “occupancy” (MacKenzie, 2005; MacKenzie et al., 2006; Latif et al., 2016).

The term “use” prevents misleading claims about occupancy, but it does not resolve the problem of distinguishing the three fundamentally different components of the detection process (described above) when conducting point counts or call-response surveys. Only a concurrent use of radio-telemetry or other methods with detection probability approaching 1.0 at the relevant scale can disentangle the probability of use from the probability of detection. MacKenzie et al. (2006, 105-106) suggest that random movements in and out of the site, often described as temporary emigration, should not bias the occupancy estimator. They drew on Kendall’s (Kendall, 1999) results regarding closure violations when estimating population size with capture-mark-recapture models. However, few studies using occupancy modelling have attempted to determine empirically the extent of temporary emigration during surveys, as well as the potential contribution to nondetection from cryptic behaviour of individuals present and their impact on occupancy estimation (Betts et al., 2008; Watson et al., 2008; Rota et al., 2009).

In the present study, our objective was to evaluate empirically the implications of temporary emigration on estimates of occupancy, using the Spruce Grouse (*Falcipennis canadensis*, hereafter, ‘grouse’) as a model species. Grouse have relatively large home ranges (16 ha; Ross, 2007), they commonly occur throughout Canadian boreal forests, but are scarce at the southern limit of their range (Whitcomb et al., 1996). Previous attempts at single-season occupancy models revealed a low detection probability of the species, possibly due to temporary emigration (Lycke et al., 2011). Here, we estimate the probability of detecting radio-marked spruce grouse given presence at the sampling site (the product *p_a_ p_d_*) and distinguish it empirically from the probability of presence at the site (*ψ*). We examine the effect of grouse movements during the survey on the detection process. Thus, we focus on detection at the individual level rather than the species level, as is typical of occupancy studies on individuals with large home ranges (Pozzanghera et al., 2016; Saracco et al., 2011). We also provide a direct comparison of an occupancy model confounding *p_a_* and *p_d_* vs. an occupancy model estimating grouse detection probability *p* strictly as *p_d_*.

## Methods

We conducted grouse call-response surveys in 2010, in the lowlands of the Saint Lawrence River, in southern Quebec, Canada (46° 30’N, 71° 21’W). We established sampling sites in coniferous stands within a landscape of 184 km^2^ composed of semiforested ombrotrophic peatlands, agricultural lands, and small towns. Coniferous forests were mostly dominated by black spruce (*Picea mariana*) mixed with eastern larch (*Larix laricina*), balsam fir (*Abies balsamea*), and eastern white cedar (*Thuja occidentalis*). The understory on these poorly drained soils was characterized by ericaceous shrubs such as blueberries (*Vaccinium* spp.), Labrador tea (*Rhododendron groenlandicum*), leatherleaf (*Chamaedaphne calyculata*), and sheep laurel (*Kalmia angustifolia*). Ericaceous shrubs were accompanied by mountain holly (*Ilex mucronata*), Appalachian tea (*Viburnum cassinoides*), and a ground layer of *Sphagnum* mosses.

### Field procedures

#### Capture and marking

We captured grouse using noose poles, either in spring by attracting them with playbacks of female territorial calls or in winter flocks (Zwickel and Bendell, 1967). Grouse were fitted with a 17 g radio transmitter (A1260 model; Advanced Telemetry Systems, Inc., Isanti, MN, USA) with a Kevlar backpack harness, individually marked with a combination of numbered plastic color bands, and released at the site of capture. Handling was authorized by Animal Care Certificate no. 08-00-05, Government of Québec, Ministère des Forêts, de la Faune et des Parcs.

#### Detection trials

In 2010, two observers conducted call-response surveys (Buckland et al., 2006) on 13 radio-marked females and 11 radio-marked males. Surveys were done at least one day after capture and release of the birds. The call-response observers broadcasted recordings of female territorial calls during 15 minutes, using 5W portable amplifiers. All grouse seen or heard were recorded according to sex with no distance limit (Schroeder and Boag, 1989). In the case of visual detections, the observers recorded the time of detection and the location of the bird using a GPS receiver, as well as its identity based on coloured leg bands. A third observer recorded the telemetry locations of a given individual, at the beginning and at the end of the survey, by walking to those locations, and recording the position with a GPS. These GPS locations enabled us to measure the distance between the grouse and the observer at the beginning and at the end of each survey. We conducted 1-5 detection trials on each grouse (*n* = 2, 3, 9, 4, and 6 individuals, respectively), with 8.5 ± 6.8 days between trials to avoid habituation to territorial call playbacks. To maximize detections, we located detection trials at random points within 200 m of the most recent grouse location prior to the trial. We conducted 81 detection trials between 22 April and 27 May 2010, between 7:30 and 16:00 EST, on days with no precipitation and wind speed < 25 km · hr^-1^.

To account for heterogeneity in detection probability among sites, we measured lateral visual obstruction at 15 m from the observer, on both sides of each sampling point on a north-south axis, using a coverboard (2.0 m height x 0.3 m width) divided into four 50 x 30 cm squares (Nudds, 1977). We expressed visual obstruction as the proportion of squares covered partially or completely.

### Statistical analysis

#### Grouse behaviour during surveys

We evaluated availability for detection in two ways. First, based on GPS locations obtained by telemetry, we calculated the distance between the observers and the grouse at the beginning and at the end of each detection trial, as well as the distance between initial and final grouse positions. Second, we classified responses to playback during the detection trial as “withdrew” (the grouse moved > 3 m away from the observers), “approached” (moved > 3 m towards the observers), “sideways” (moved but remained at the same distance to the observers), and “stationary”. We focused on these three movement responses to playback because we expected that grouse mobility during a survey influences detection probability of the individual.

#### Estimating components of detection

We defined sampling sites spatially as the circular areas centered on the call-response observers, with a radius corresponding to the maximum distance at which grouse were detected by radio-telemetry observers. Thus, grouse were deemed “present” when they were within the maximum detection distance. Birds were deemed “available for detection” if they were present within the maximum detection distance and were not stationary during the survey. We compared detection rates in relation to the three movement responses with Fisher’s exact test. Besides presence within sampling sites, we modelled other possible influences on detection with generalized linear mixed models (GLMMs) (Bates et al., 2015), with a binomial response variable and grouse ID as a random effect. These possible influences on detection during surveys included the nearest distance of grouse to call-response observers, sex, Julian day, minutes since sunrise, and mean percent lateral cover.

We compared detection probabilities calculated directly as proportions of “present” or “available” birds that were observed *vs*. estimates obtained from single-season site-occupancy models (MacKenzie et al., 2002) implemented with the R package ‘unmarked’ (Fiske et al., 2011; R Core Team, 2020). For a given individual, surveys were conducted at different sites. Thus, surveys did not provide the repeated visits to the same sites required for occupancy analysis. To circumvent this issue, we used a simulation approach to investigate the potential influence of temporary emigration and availability for detection on occupancy and detection probability estimates. We proceeded by six steps. 1) we used the first survey site of each individual, and calculated distances between grouse location at the time of each real survey and the center of the original call-response site. In cases where several grouse locations were georeferenced during a single survey, we used the mean of location coordinates. 2) we inserted the distances calculated in the first step into the linear predictor of the GLMM described above to obtain predicted probabilities. 3) we performed Bernoulli trials with *p* parameters corresponding to predicted probabilities obtained in the second step. 4) we organized the 1’s and 0’s obtained in step three into detection histories for each original site. All visits were included in the simulation; thus, detection histories were simulated as if observers had conducted all surveys in the original set of call-response sites. 5) we estimated by maximum likelohood *ψ* and *p* from an intercept-only occupancy model, i.e. with detection probability and occupancy held constant. Finally, we replicated steps 3) to 5) 1000 times and report means, 2.5^th^ and 97.5^th^ percentiles of the empirical distribution of *ψ* and *p*. Note that all sites used in the simulation were known to be occupied; hence the only random process simulated here was the imperfect detection. We conducted the simulation exercise for three different scenarios: 1) all surveys, 2) surveys with only grouse known to be within the maximum detection distance, and 3) surveys with only grouse present and available for detection (as determined empirically from their behaviour).

## Results

We detected grouse in 29.6 % of the 81 detection trials (95 % confidence interval [0.20, 0.41]). The maximum detection distance from observers was 83 m, thus sampling sites were considered as 83 m radius circles centered on the observer. Grouse moved from their initial location during 72.8 % of the detection trials, with a median distance of 12.3 m. Twenty-nine percent of the grouse that moved approached the observers, 23 % moved sideways, and 42 % withdrew relative to the observers. The observers detected 68 %, 27 %, and 25 % of the grouse that approached, moved sideways, or withdrew (Fisher’s exact test, p < 0.009). In accordance to our definition of availability for detection (see Methods), we detected none of the six stationary grouse that were present at the site during the surveys. Besides their movements (or lack thereof) during the survey, only the distance of the grouse significantly affected the detection probability (Figure 1).

**Figure 1:**
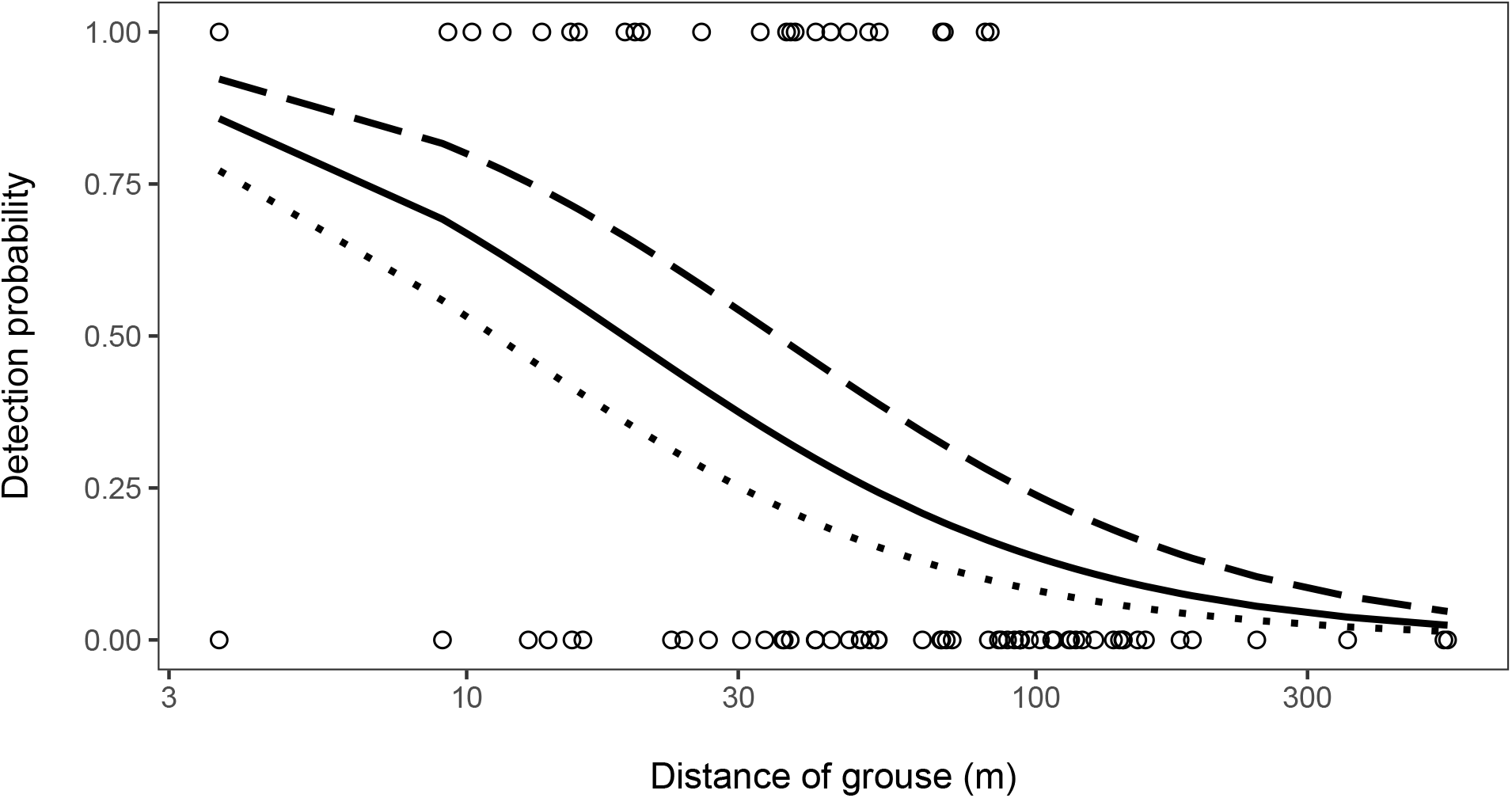
Relationship between Spruce Grouse detection probability, and distance to observers during 15-min call-response surveys. Each dot represents one survey. Dotted, continuous, and dashed lines represent generalized linear model estimates at the earliest, median, and latest sampling dates, respectively. Data collected in spring 2010 based on point counts and presence confirmed from radio-telemetry in southern Quebec, Canada.

Sex, lateral cover, time of day, and Julian date had no significant effect on grouse detection (Table 1). Grouse were present at the site, i.e., inside the detection radius during 53 of the 81 detection trials (*p_p_* = 0.65 [0.54-0.76]). They would have been present in 34 (42 %) of the trials ([0.31-0.53]), had we remained at the site of the first visit of each individual throughout the study. The *p_p_* estimate was significantly lower than the corresponding occupancy estimate obtained from simulation (*ψ* = 0.89, [0.61, 1]). Forty-six of the 53 grouse present at the site were available for detection (i.e., moved and became conspicuous during the survey; *p_a_* = 0.89 [0.77, 0.96]). Twenty-four of the 47 available grouse were detected (*p_d_* = 0.51 [0.36, 0.66]). Thus, the detection probability of grouse given presence (*p_a_p_d_*) was 0.45 (24/53, [0.32, 0.60]), i.e. twice the corresponding detection probability obtained from the mean estimates of the 1000 occupancy model simulations (*p* = 0.23, [0.12, 0.35]). When grouse absent from the site during the survey were dropped, occupancy and its precision remained similar (*ψ* = 0.88, [0.57, 1]), but detection probability became very similar to the direct measurement (p = 0.42 [0.19, 0.65]).

**Table 1:**
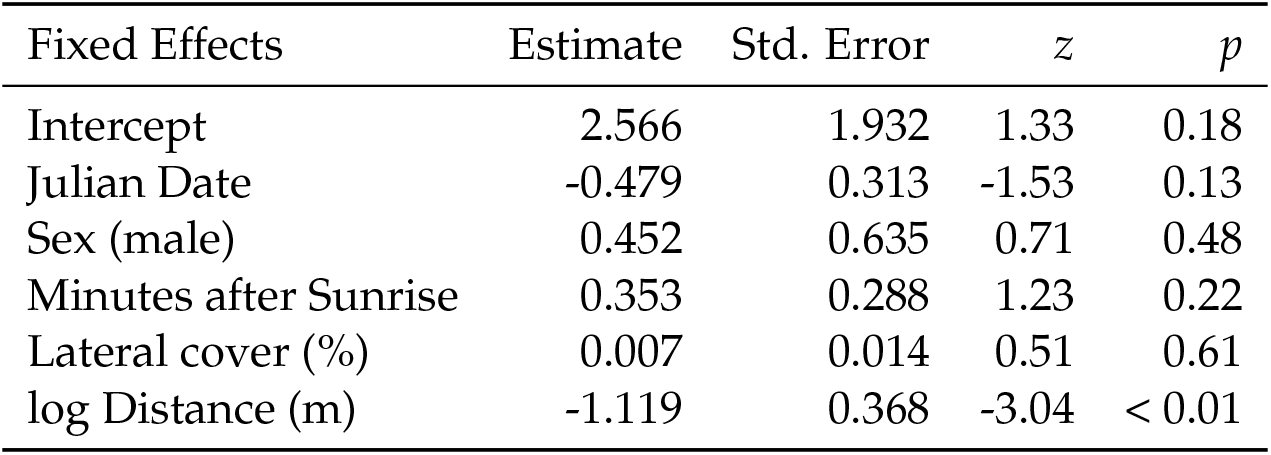
Generalized linear mixed model influences of Spruce Grouse detection probabilities, as determined by a generalized linear mixed model with Site ID as a random effect. Residual deviance = 79.5, residual degrees of freedom = 74. Data collected in spring 2010 based on point counts and presence confirmed from radio-telemetry in southern Quebec, Canada.

## Discussion

Based on a novel combination of call-response surveys and simultaneous radio-tracking, we showed that occupancy estimates can be robust to deviations from major assumptions about the components of the detection process, i.e., temporary emigration, availability to be detected, and detection of ‘available’ birds by the observers. However, estimates of detection probability by occupancy models were highly sensitive to temporary emigration.

Single-season occupancy models assume closure, i.e., the absence of change in the occupancy state at a site throughout the sampling period (MacKenzie et al., 2002). However, birds often make extraterritorial forays during the breeding season (Whitaker and Warkentin, 2010) and these forays can change the state of the site, especially when individuals occur at very low densities. Rota et al. (2009) attributed changes in site occupancy estimates to closure violations, mainly caused by local colonization and extinction over the sampling period. Watson et al. (2008) and McClure and Hill (2012) also reported temporary emigration in bird species, highlighting that dynamic occupancy models performed better than single season occupancy models with their data (Watson et al., 2008). Otto et al. (2013) observed changes in the occupancy of terrestrial salamanders within the season and noted a positive bias of occupancy in response to closure violations. In contrast, ignoring temporary emigration by grouse in our study did not influence occupancy estimates, but resulted in underestimating detection probability *sensu stricto*. Results therefore vary substantially among the few empirical assessments of the consequences of temporary emigration on occupancy estimates.

Occupancy models can be influenced by the ratio of home range size to sampling plot size (Efford and Dawson, 2012). Spruce Grouse have very large annual home ranges (16 ha; Ross 2007) relative to the size of sampling plots in which birds can be detected. However, grouse home ranges in the spring may become much smaller, e.g. 1.2 – 8.5 ha in April-May (Ellison, 1973). The magnitude of temporary emigration and resulting bias in occupancy likely differ among lifehistory traits and habitat types. If birds modify their home range size in response to changes in habitat (e.g., local breeding density, landscape composition, lateral vegetation cover, among others), resulting changes in temporary emigration may affect parameter estimates and inferences on the drivers of species’ distributions. Birds can enlarge their home range to acquire sufficient resources in response to habitat loss and fragmentation caused by forest harvesting (Gjerde and Wegge, 1989; Leonard et al., 2008). For instance, Norris and Stutchbury (2001) showed that male hooded warblers (*Setophaga citrina*) moved more extensively in the breeding season and spent more time off their territories in fragmented habitat compared to continuous habitat, presumably because of lower availability of neighbouring extra-pair mates.

The consequences of bias in occupancy model estimates could be severe for declining populations in patchy habitats that are of particular concern for management and conservation, as suggested by Rota et al. (2009). For example, differences in the probability of temporary emigration due to variation in home range size among habitats or across time could be interpreted as differences in occupancy over space or time.

Temporary emigration may be addressed in various ways in studies focusing on occupancy with various modifications of Pollock’s robust design (Pollock, 1982). MacKenzie et al. (2003) used multiple surveys within the season (i.e., primary sampling periods), and partitioned them into shorter sampling intervals (i.e., secondary sampling periods). Moreover, under spatial subsampling with replacement within a sampling unit, multi-scale occupancy models provide a means of estimating temporary emigration in the spatial sub-units, given that the species occupies the sampling unit. The closure assumption can also be violated within primary periods (Norris and Stutchbury, 2001). Valente et al. (2017) addressed this issue in a dynamic occupancy setting, where they observed biased dynamic rates (colonization and extinction) when temporary emigration within a primary period. Finally, accounting for temporary emigration also extends to investigations of occupancy patterns in species assemblages (Kéry et al., 2009).

### Management Implications

We showed that a substantial underestimation of detection probability may arise in spruce grouse surveys because of temporary emigration. Detection probability, even with grouse present, was low. Thus, there is scope for substantial improvement in sampling efficiency, for example with changes in the field protocol that would induce grouse to move, especially nearer the observer. Here, occupancy estimates remained robust to this violation of the closure assumption, but this conclusion may not apply to other systems.

A tempting solution to alleviate problems related to the closure assumption would be to enlarge the area of sampling sites. However, this solution rapidly reaches a limit within a call-response survey design if only a single point is surveyed at each site. In this design, the observer remains stationary, in which case sampling beyond the detection range (e.g. 83 m in this study) would be pointless, because individuals beyond this range are not detectable. A more viable solution is to conduct multiple call-response surveys at different points or transects deployed within larger sites (e.g., an order of magnitude larger than species home range). This would simultaneously increase detection probability and alleviate the problem of temporary emigration, because more than a single individual is likely to occupy the site at any given moment. For example, using transects in New York State, Ross et al. (2016) reached a detection probability of 0.65. This multiple-sampling point approach has at least two advantages: it increases the number of individuals within detection range, and the movements of the observers may induce movements by the targeted individuals, which can increase the detection probability, as shown in the present study.

## Acknowledgments

We are grateful to Céline Macabiau for her major contribution to field work and data management. We thank S. St-Onge, A. Desrosiers, P. Beaupré, B. Baillargeon, D. Grenier, C. Daigle, and J. Fortin for captures and telemetry monitoring of grouse. We thank M.-P. Amyot, C. Descamps, C. Lagacé, M. Paquet, and G. Sauvestre for field assistance in the detection trials. Thanks also go to Solifor Inc. and the many landowners for permission to conduct fieldwork on their properties. We thank W. F. J. Parsons for a thorough review of the English of the revised version. Financial support for this study was provided by the Ministère des Forêts, de la Faune et des Parcs du Québec (MFFP), the Fondation de la faune du Québec, and the Natural Sciences and Engineering Research Council of Canada (NSERC) – Laval University Industrial Research Chair in Peatland Management.

## References

Bates, D., M. Maechler, B. Bolker, and S. Walker. 2015. Fitting Linear Mixed-Effects Models Using lme4. Journal of Statistical Software 67:1–48.

Betts, M. G., N. L. Rodenhouse, T. S. Sillett, P. J. Doran, and R. T. Holmes. 2008. Dynamic occupancy models reveal within-breeding season movement up a habitat quality gradient by a migratory songbird. Ecography 31:592–600.

Buckland, S. T., R. W. Summers, D. L. Borchers, and L. Thomas. 2006. Point transect sampling with traps or lures. Journal of Applied Ecology 43:377–384.

Efford, M. G. and D. K. Dawson. 2012. Occupancy in continuous habitat. Ecosphere 3:1–15.

Ellison, L. N. 1973. Seasonal social organization and movements of Spruce Grouse. The Condor 75:375–385.

Fiske, I., R. B. Chandler, and A. Royle. 2011. Unmarked: Models for data from unmarked animals. R package version 0.11-0. http://CRAN.R-project.org/package=unmarked.

Gjerde, I. and P. Wegge. 1989. Spacing pattern, habitat use and survival of capercaillie in a fragmented winter habitat. Ornis Scandinavica 20:219–225.

Kendall, W. L. 1999. Robustness of closed capture-recapture methods to violations of the closure assumption. Ecology 80:2517–2525.

Kéry, M., J. A. Royle, M. Plattner, and R. M. Dorazio. 2009. Species richness and occupancy estimation in communities subject to temporary emigration. Ecology 90:1279–1290.

Kéry, M. and B. R. Schmidt. 2008. Imperfect detection and its consequences for monitoring for conservation. Community Ecology 9:207–216.

Latif, Q. S., M. M. Ellis, and C. L. Amundson. 2016. A broader definition of occupancy: comment on Hayes and Monfils. Journal of Wildlife Management 80:192–194.

Leonard, T. D., P. D. Taylor, and I. G. Warkentin. 2008. Landscape structure and spatial scale affect space use by songbirds in naturally patchy and harvested boreal forests. The Condor 110:467–481.

Lycke, A., L. Imbeau, and P. Drapeau. 2011. Effects of commercial thinning on site occupancy and habitat use by spruce grouse in boreal Quebec. Canadian Journal of Forest Research 41:501–508.

MacKenzie, D. I. 2005. Was it there? Dealing with imperfect detection for species presence/absence data. Australian & New Zealand Journal of Statistics 47:65–74.

MacKenzie, D. I., J. D. Nichols, J. E. Hines, M. G. Knutson, and A. B. Franklin. 2003. Estimating site occupancy, colonization, and local extinction when a species is detected imperfectly. Ecology 84:2200–2207.

MacKenzie, D. I., J. D. Nichols, S. Lachman, S. Droege, A. Royle, and C. A. Langtimm. 2002. Estimating site occupancy rates when detection probabilities are less than one. Ecology 83:2248–2255.

MacKenzie, D. I., J. D. Nichols, J. A. Royle, K. H. Pollock, L. L. Bailey, and J. E. Hines. 2006. Occupancy estimation and modeling: inferring patterns and dynamics of species occurrence. Academic Press, New York, NY, USA.

Mazerolle, M. J., A. Desrochers, and L. Rochefort. 2005. Landscape characteristics influence pond occupancy by frogs after accounting for detectability. Ecological Applications 15:824–834.

McClure, C. J. W. and G. E. Hill. 2012. Dynamic versus static occupancy: how stable are habitat associations through a breeding season? Ecosphere 3:60.

Norris, D. R. and B. J. M. Stutchbury. 2001. Extraterritorial movements of a forest songbird in a fragmented landscape. Conservation Biology 15:729–736.

Nudds, T. D. 1977. Quantifying the vegetative structure of wildlife cover. Wildlife Society Bulletin 5:113–117.

Otto, C. R. V., L. L. Bailey, and G. J. Roloff. 2013. Improving species occupancy estimation when sampling violates the closure assumption. Ecography 36:1299–1309.

Pollock, K. H. 1982. A capture-recapture design robust to unequal probability of capture. Journal of Wildlife Management 46:752–757.

Pozzanghera, C. B., K. J. Sivy, M. S. Lindberg, and L. R. Prugh. 2016. Variable effects of snow conditions across boreal mesocarnivore species. Canadian Journal of Zoology 94:697–705.

R Core Team. 2020. R: A Language and Environment for Statistical Computing. R Foundation for Statistical Computing, Vienna, Austria.

Ross, A. M. 2007. Spruce grouse distribution, movements and habitat selection: a mid-successional species in an aging forested landscape. Ph.D. thesis, State University of New York.

Ross, A. M., G. Johnson, and J. Gibbs. 2016. Spruce Grouse decline in maturing lowland boreal forests of New York. Forest Ecology and Management 359.

Rota, C., J. Fletcher, R. J., R. M. Dorazio, and M. G. Betts. 2009. Occupancy estimation and the closure assumption. Journal of Applied Ecology 46:1173–1181.

Saracco, J. F., R. B. Siegel, and R. L. Wilkerson. 2011. Occupancy modeling of black-backed woodpeckers on burned Sierra Nevada forests. Ecosphere 2:31.

Schroeder, M. A. and D. A. Boag. 1989. Evaluation of a density index for territorial male spruce grouse. Journal of Wildlife Management 53:475–478.

Valente, J. J., R. A. Hutchinson, and M. G. Betts. 2017. Distinguishing distribution dynamics from temporary emigration using dynamic occupancy models. Methods in Ecology and Evolution 8:1707–1716.

Watson, C., F. Weckerly, J. Hatfield, C. Farquhar, and P. Williamson. 2008. Presencenonpresence surveys of goldencheeked warblers: detection, occupancy and survey effort. Animal Conservation 11:484–492.

Whitaker, D. M. and I. G. Warkentin. 2010. Spatial ecology of migratory passerines on temperate and boreal forest breeding grounds. The Auk 127:471–484.

Whitcomb, S. D., F. A. Servello, and J. O’Connell, A. F. 1996. Patch occupancy and dispersal of Spruce Grouse on the edge of its range in Maine. Canadian Journal of Zoology 74:1951–1955.

Zwickel, F. C. and J. F. Bendell. 1967. A snare for capturing blue grouse. Journal of Wildlife Management 31:202–204.

